# MatchCLOT: Single-Cell Modality Matching with Contrastive Learning and Optimal Transport

**DOI:** 10.1101/2022.11.16.516751

**Authors:** Federico Gossi, Pushpak Pati, Adriano Martinelli, Maria Anna Rapsomaniki

## Abstract

Recent advances in single-cell technologies have enabled the simultaneous quantification of multiple biomolecules in the same cell, opening new avenues for understanding cellular complexity and heterogeneity. However, the resulting multimodal single-cell datasets present unique challenges arising from the high dimensionality of the data and the multiple sources of acquisition noise. In this work, we propose MatchCLOT, a novel method for single-cell data integration based on ideas borrowed from contrastive learning, optimal transport, and transductive learning. In particular, we use contrastive learning to learn a common representation between two modalities and apply entropic optimal transport as an approximate maximum weight bipartite matching algorithm. Our model obtains state-of-the-art performance in the modality matching task from the NeurIPS 2021 multimodal single-cell data integration challenge, improving the previous best competition score by 28.9%. Our code can be accessed at https://github.com/AI4SCR/MatchCLOT.

## 1 Introduction

Single cells are complex dynamical systems where a variety of biomolecules interact in a coordinated way to produce robust and adaptive behaviors. Understanding the causal relationships between these biomolecules and their role in health and disease is a longstanding question in biology and medicine [10]. Recent advances in single-cell technologies have made it possible to simultaneously quantify different combinations of (epi)genomic, transcriptomic, and proteomic profiles [19]. Integrated analysis of the resulting multiomic profiles has the potential to capture cellular state, complexity and heterogeneity at an unprecedented scale. However, a number of emerging challenges, associated with the high dimensionality and acquisition noise in the resulting datasets, limit this potential [8]. To address these challenges, the single-cell community launched a multimodal single-cell data integration challenge at NeurIPS 2021 [12]. The challenge provided a curated multimodal benchmarking dataset and defined three tasks, namely, modality prediction, modality matching, and joint representation across modalities. Here we propose MatchCLOT (Single-cell modality MATCHing with Contrastive Learning and Optimal Transport), a novel solution that addresses modality matching, *i*.*e*., predicting the cell matching between two sets of single-cell profiles from different omic modalities. Our work is inspired by recent promising applications of optimal transport (OT) in various single-cell data analysis tasks ([1, 3, 7, 9, 14, 18, 20]). MatchCLOT is built on top of CLIP [17], a contrastive learning model also used by Team Novel in the challenge [12]. Unlike Team Novel’s model that uses a maximum weight bipartite graph and generates hard matchings, we propose a novel OT solution that allows MatchCLOT to generate soft matchings, leading to increased performance and significant gains in computational time and memory. MatchCLOT additionally exploits prior knowledge of the batch label, resulting in a smaller search space, and uses a transductive learning setup that mitigates the effects of distribution shifts in the test data. Overall, MatchCLOT obtains the state-of-the-art result, improving over the competing best method, scMoGNN [21], by 28.9% for the *overall matching probability score* and by 209% for a *top-5 matching accuracy* metric.

## 2 Methods

MatchCLOT consists of three different modules, shown in different colors (Figure 1). *Preprocessing* includes normalization, low-dimensional projection and batch effect correction in a transductive setting. *Training* leverages a contrastive learning setup to maximize the similarity between matching cell profiles in the embedding space. Finally, *testing* involves an entropic regularized differentiable OT and utilization of batch labels for predicting the matching cell profiles.

**Figure 1:**
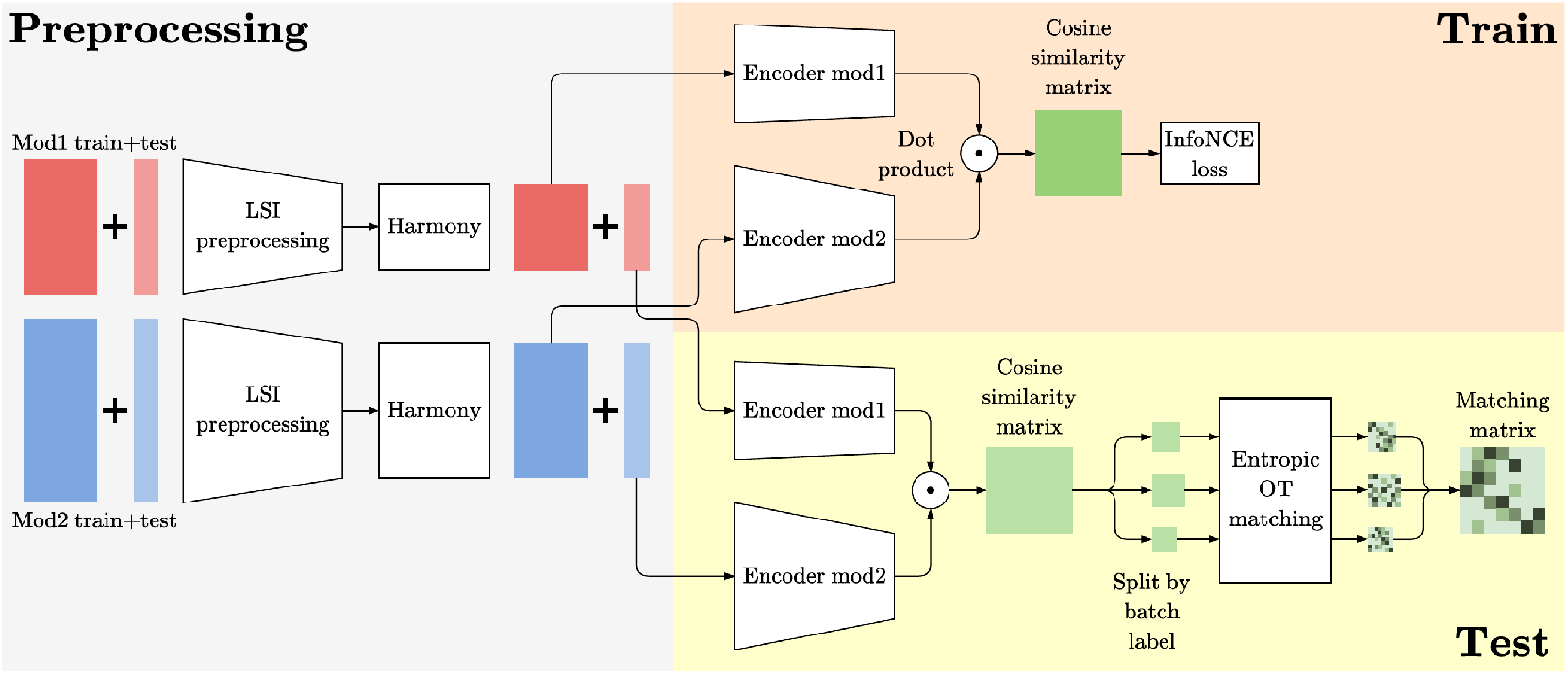
Overview of our proposed MatchCLOT framework.

### Preprocessing

The first component of MatchCLOT preprocesses the raw single-cell data in a transductive setting, by operating on the union of the train and test set that is available while training. For each modality, MatchCLOT normalizes and reduces the dimensions of the combined train and test set using latent semantic indexing (LSI), a common method for processing scATAC-seq data [5]. LSI consists of a term frequency-inverse document frequency (TF-IDF) normalization coupled to an L1-normalization and a log(1 + *x*) transformation, and followed by truncated SVD and a zero mean and unit variance scaling. LSI results in preprocessed measurements of lower dimensions, that are in turn corrected for batch effects using Harmony [11], an established batch effect correction method. Since the test data batches are independent of the training data batches, correcting the batch effects in a transductive setting minimizes the impact of the distribution shifts due to acquisition variations in the data. We note that during the transductive preprocessing, the two modalities are processed independently and the ground-truth labels are not used by the model. We also note that leveraging the unlabeled test data was a common practice during the competition, as several methods applied unsupervised approaches on the test data prior to model training.

### Contrastive learning and Encoder architecture

The backbone of MatchCLOT is built on CLIP [17], a contrastive learning model, aiming to generate similar latent representations for profiles of the same cell and orthogonal representations for non-matching profiles. Both encoders, as shown in Figure 1, are shallow multi-layer perceptrons (MLPs) designed as per the modalities. The output embedding dimensions of the encoders are also set according to the operating subtask, *i*.*e*., CITE or Multiome. Modality-specific embeddings are unit normalized and multiplied via a dot product to produce a cosine similarity matrix. Afterwards, an InfoNCE [15] contrastive objective is calculated for all the embedding pairs in the similarity matrix for backpropagation. We tune the model and training hyperparameters, namely LSI reduced dimensions, encoder hidden layer dimensions, dropout rate, learning rate, and weight decay, on a validation split obtained from the labeled training data.

### Optimal transport matching

The method proposed by Team Novel used a maximum weight bipartite perfect matching to generate a matching prediction from the cosine similarity matrix of the multiomic profiles. Instead, we propose to use OT for faster and better matching. OT is a field of mathematics that studies the best way of transporting a source distribution to a target distribution while minimizing the costs of displacement. Given two discrete probability distributions with supports *A, B* of the same size (|*A*| = |*B*| = *n*), with densities *α, β* and costs *c*(*a, b*) defined ∀*a* ∈ *A*, ∀*b* ∈ *B*, the linear program formulation of the OT problem is given as,

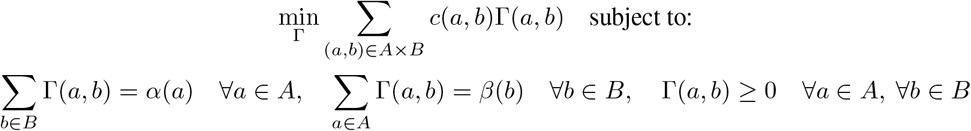

To use OT as an approximation of the max-weight bipartite matching, we can relax the integer linear program (ILP) formulation of the max-weight bipartite matching and convert it to an OT problem. Given a bipartite graph *G* = (*V, E*) with bipartition (*A, B*), weight function *w* : *E* ↦ ℝ, a matching *M* ⊆ *E*, let *x*(*a, b*) = 1, if (*a, b*) ∈ *M* and 0 otherwise. Then, the ILP formulation of the maximum weight bipartite perfect matching is as follows,

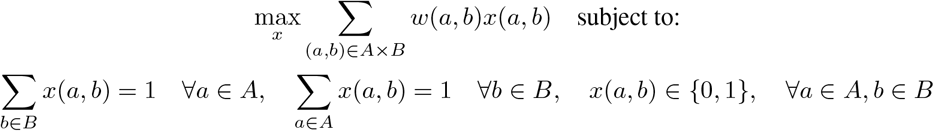

By dropping the integrality constraint on the variables *x*(*a, b*), the problem becomes an LP and can be converted to an OT problem. The corresponding OT problem uses negative weights −*w*(*a, b*) as costs *c*(*a, b*), a function of variables 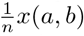 as transport plan Γ(*a, b*), and uniform distributions over *A, B* with densities 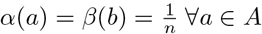, ∀*b* ∈ *B*. With the OT formulation, the transport plan Γ can be interpreted as a soft matching, where each vertex *a* ∈ *A* can be matched with multiple vertices *b* ∈ *B*. Adding an entropic regularization [6] can speed up the computation of OT and leads to the following objective,

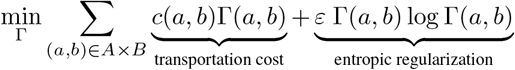

The term *ε* determines the strength of the entropic regularization, with higher values producing noisier transport plans Γ. In our case, we use *ε* = 0.01 to generate a soft matching from the cosine similarity matrix.

### Matching with batch label information

To avoid matching cell profiles from different batches, MatchCLOT exploits the test data batch labels to reduce the search space for the matching algorithm. This is achieved by splitting the profiles by batch labels, computing the cosine similarity matrices and entropic OT matching per batch, and combining the matching matrices for the final prediction. We note that the batch labels were available to all teams during the competition, and were also exploited by some of the top-scoring methods (e.g., Team CLUE and scMoGNN).

## 3 Results and Discussion

### Dataset

We benchmarked MatchCLOT on the modality matching task from NeurIPS 2021 multimodal single-cell data integration challenge [13]. The dataset processed bone marrow samples from 12 donors at 4 data generation sites via two multiomic single cell technologies: CITE-seq, which captures single-cell RNA gene expression (GEX) and surface protein levels (as Antibody Derived Tags, ADT); and the 10X Multiome assay, which captures chromatin accessibility (based on the Assay for Transposase-Accessible Chromatin, ATAC) and single-nucleus RNA gene expression (GEX) levels. The CITE-seq and Multiome data included 90,000 and 70,000 cells, respectively. The test data consists of 15,066 cells for the CITE subtasks GEX2ADT and ADT2GEX, and 20,009 cells for the Multiome subtasks ATAC2GEX and GEX2ATAC.

### Evaluation metrics

The NeurIPS challenge measures the modality matching performance of a method in terms of the weight/probability assigned to the correct cell pairings. Provided a probability matching matrix **M** ∈ ℝ^*n*×*n*^ for *n* cells, the *matching probability score* is computed as,

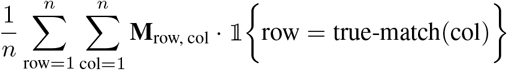

Additionally, we measure the *top-K matching accuracy*, which quantifies the accuracy of matching the top score in the rows/columns of **M** with the correct cell pairings, given as,

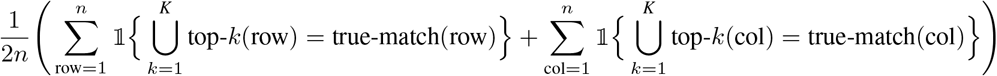

### Implementation

We implemented MatchCLOT using PyTorch [16] and conducted the experiments on NVIDIA Tesla P100 GPU and POWER9 CPU. The labeled CITE-seq and Multiome data was split into two sets according to the batch labels for method validation (1 batch) and training (8-9 batches). The model and training hyperparameters were optimized using Wandb [2], a bayesian hyperparameter optimization library. The comprehensive list of hyperparameters are provided in the Appendix Tab. 2.

### Quantitative analysis

The matching probability scores of MatchCLOT and the competing methods for five modality matching subtasks, *i*.*e*., GEX2ATAC, ATAC2GEX, ADT2GEX, GEX2ADT, and Overall, is presented in Table 1. Team CLUE and Team Novel ranked first and second, respectively, among a total of 462 submissions from 23 teams, across all the subtasks in the NeurIPS challenge. scMoGNN [21] is a post-challenge method that outperformed the winners. MatchCLOT achieved the state-of-the-art scores for all the subtasks and improves over scMoGNN by 28.9% for the overall matching score. For the top-5 matching accuracy, MatchCLOT produced 0.2226 compared to 0.0720 by scMoGNN, an improvement of 209%. We note that, as scMoGNN follows a hard-matching approach, the competition score and the top-5 matching accuracy of their predictions are identical. This significant gain demonstrates the superiority of the matching matrix **M**, where the paired cellular profiles appear among the top-5 probability scores across the rows and the columns. Noticeably, MatchCLOT is independent of the sequence of the modality matching task, *i*.*e*., GEX2ATAC & ATAC2GEX, and ADT2GEX & GEX2ADT are the same, similar to Team Novel and scMoGNN.

**Table 1:**
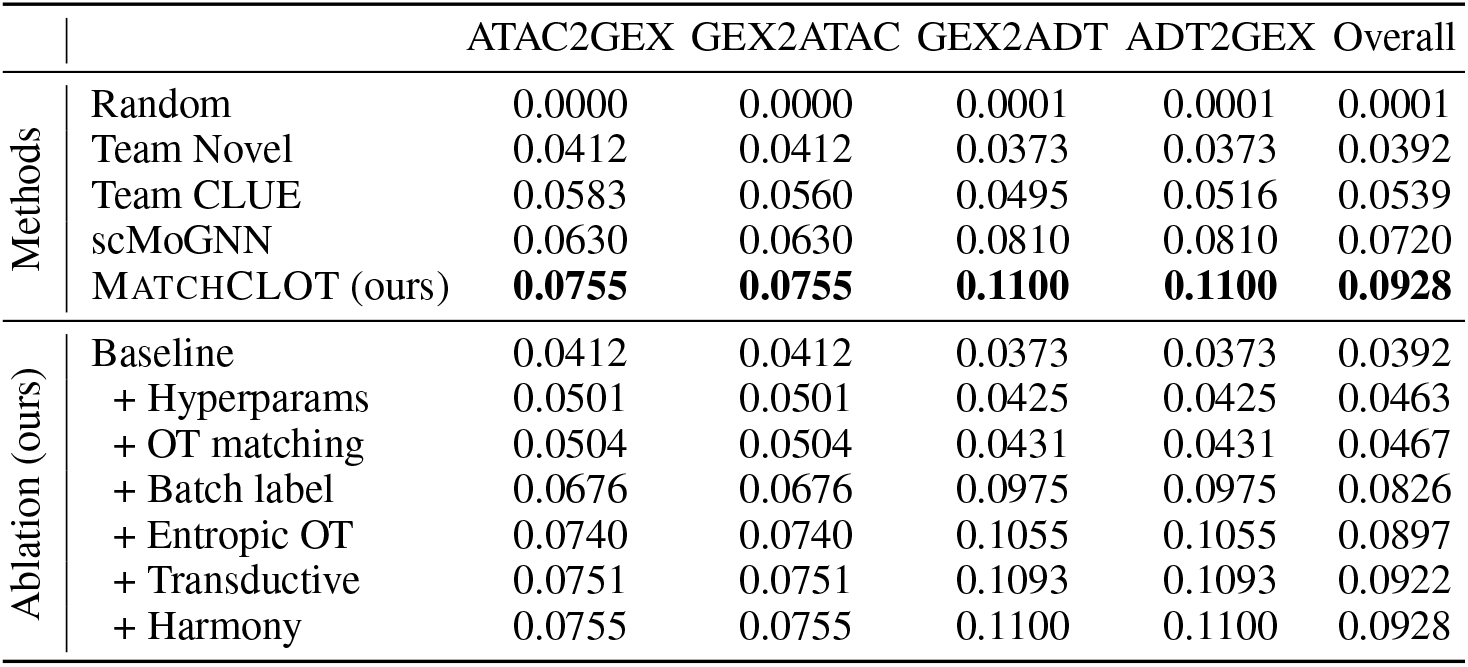
Cell-level modality matching scores for competing methods. Task-wise best results in **bold**.

|  |  | ATAC2GEX | GEX2ATAC | GEX2ADT | ADT2GEX | Overall |
| --- | --- | --- | --- | --- | --- | --- |
| Methods | Random | 0.0000 | 0.0000 | 0.0001 | 0.0001 | 0.0001 |
|  | Team Novel | 0.0412 | 0.0412 | 0.0373 | 0.0373 | 0.0392 |
|  | Team CLUE | 0.0583 | 0.0560 | 0.0495 | 0.0516 | 0.0539 |
|  | scMoGNN | 0.0630 | 0.0630 | 0.0810 | 0.0810 | 0.0720 |
|  | MATCHCLOT (ours) | **0.0755** | **0.0755** | **0.1100** | **0.1100** | **0.0928** |
| Ablation (ours) | Baseline | 0.0412 | 0.0412 | 0.0373 | 0.0373 | 0.0392 |
|  | + Hyperparams | 0.0501 | 0.0501 | 0.0425 | 0.0425 | 0.0463 |
|  | + OT matching | 0.0504 | 0.0504 | 0.0431 | 0.0431 | 0.0467 |
|  | + Batch label | 0.0676 | 0.0676 | 0.0975 | 0.0975 | 0.0826 |
|  | + Entropic OT | 0.0740 | 0.0740 | 0.1055 | 0.1055 | 0.0897 |
|  | + Transductive | 0.0751 | 0.0751 | 0.1093 | 0.1093 | 0.0922 |
|  | + Harmony | 0.0755 | 0.0755 | 0.1100 | 0.1100 | 0.0928 |

Table 1 also presents the ablation studies of our method in an incremental order of different components for all the subtasks. The baseline refers to the method by Team Novel, on which MatchCLOT is built on. MatchCLOT improved over the baseline by 18.1% via thorough hyperparameter tuning. Utilizing the batch labels rendered the largest gain of 76.9%. Further gains of 8.6% and 2.8% in matching score were achieved via entropic regularization and differentiable OT, and transductive learning, respectively. Though the OT matching did not contribute to the matching score, it significantly improved the computation time and memory usage (Appendix Fig. 2). We quantified these gains on **M** ∈ ℝ^15066×15066^ over five runs for GEX2ADT subtask. Team Novel discarded 99.5% of the edges in **M** during maximum weight bipartite matching to render a computation time and memory usage of ∼125 seconds and 12GB, respectively, compared to OT using 100% of the edges within 50 seconds and 16GB. For equal memory constraint, Team Novel required to discard 95% of the edges. Although discarding small values in **M** works well in this task, in different contexts it might inhibit the performance by ignoring valuable information.

## 4 Conclusion and Future work

In this work, we proposed a solution for matching modalities across multimodal single-cell data. Our method MatchCLOT utilizes optimal transport, contrastive and transductive learning and achieves the state-of-the-art performance, while being significantly more efficient in terms of computational time and memory usage. A key challenge in our solution is to define reasonable data augmentations of single-cell omics, which is crucial for contrastive learning[4]. Future works in this direction can address this challenge, and explore the recent advances in contrastive learning techniques and investigate powerful encoding models, *e*.*g*., transformer-based encoders, to potentially improve the results along with identifying interpretable omics information utilized by the model.

## A Appendix

### Hyperparameter values

**Table 2:** Best hyperparameter configurations for the modality matching model found with bayesian optimization. The batch size was set without using bayesian optimization.

| Hyperparameter | Search space | GEX2ATAC | GEX2ADT |
| --- | --- | --- | --- |
| LSI dim mod1 | {64, 96, 128, 192, 256, 384, 512} | 192 | 192 |
| LSI dim mod2 | {64, 96, 128, 192, 256, 384, 512} | 256 | 134 |
| Encoder hidden dim mod1 | {128, 256, 512, 1024, 2048, 4096}<sup>2</sup> | (2048, 1024) | (256, 2048) |
| Encoder hidden dim mod2 | {128, 256, 512, 1024, 2048, 4096}<sup>2</sup> | (2048) | (4096, 2048) |
| Embedding dim | {128, 256, 512, 1024} | 128 | 256 |
| Dropout rates mod1 | $[0.0, 0.7]^2$ | (0.34, 0.47) | (0.3, 0.05) |
| Dropout rates mod2 | $[0.0, 0.7]^2$ | (0.67) | (0.4, 0.2) |
| Initial temperature $\log \tau$ | $[1.0, 5.0]$ | 2.74 | 4.0 |
| Learning rate | $[10^{-6}, 10^{-3}]$ | $6 \cdot 10^{-4}$ | $1.75 \cdot 10^{-4}$ |
| Weight decay | $[10^{-6}, 10^{-3}]$ | $1.25 \cdot 10^{-4}$ | $2 \cdot 10^{-4}$ |
| Batch size* | - | 16384 | 16384 |

### Efficiency benchmarking for optimal transport matching

**Figure 2:**
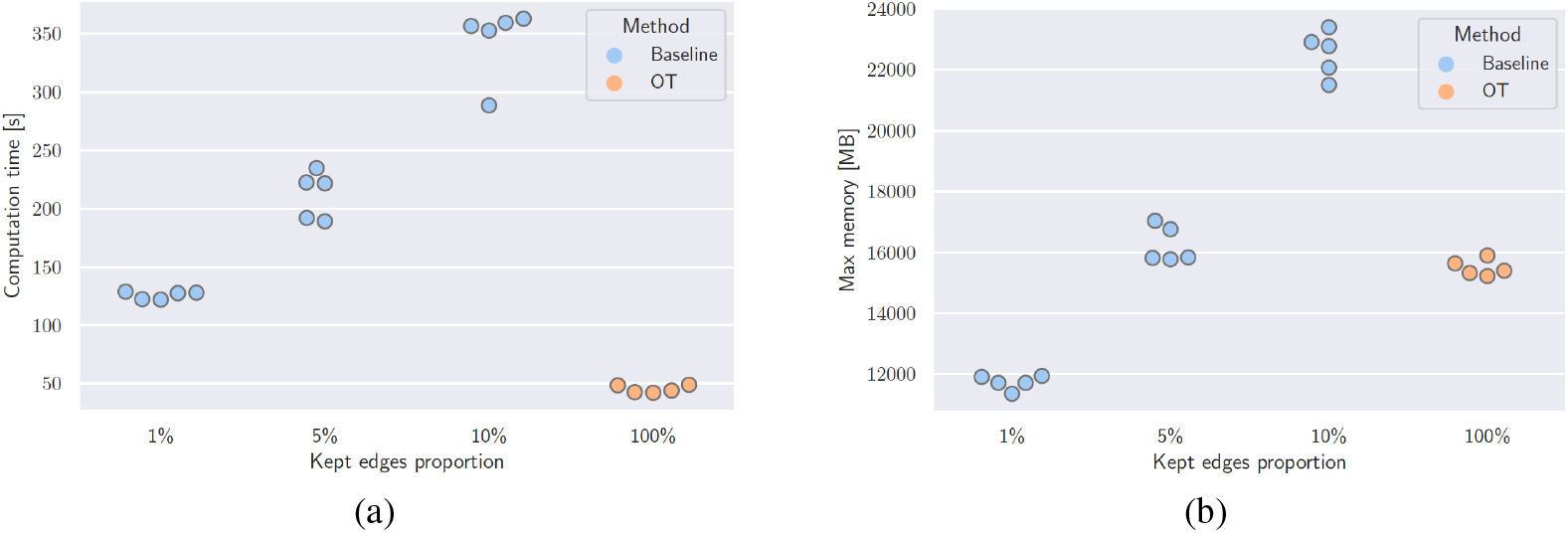
Comparison of computation time (a) and memory usage (b) between the baseline method using max-weight bipartite matching (baseline - blue) and OT matching (MatchCLOT - orange) for different proportions of retained matching edges.

## References

[1] Riccardo Bellazzi, Andrea Codegoni, Stefano Gualandi, Giovanna Nicora, and Eleonora Vercesi. The gene mover’s distance: Single-cell similarity via optimal transport. 2102.01218, 2021.

[2] Lukas Biewald. Experiment tracking with weights and biases, 2020. Software available from https://wandb.com.

[3] Charlotte Bunne, Laetitia Papaxanthos, Andreas Krause, and Marco Cuturi. Proximal optimal transport modeling of population dynamics. In International Conference on Artificial Intelligence and Statistics, pages 6511–6528. PMLR, 2022.

[4] Ting Chen, Simon Kornblith, Mohammad Norouzi, and Geoffrey Hinton. A simple framework for contrastive learning of visual representations. In International conference on machine learning, pages 1597–1607. PMLR, 2020.

[5] Darren A Cusanovich, Riza Daza, Andrew Adey, Hannah A Pliner, Lena Christiansen, Kevin L Gunderson, Frank J Steemers, Cole Trapnell, and Jay Shendure. Multiplex single-cell profiling of chromatin accessibility by combinatorial cellular indexing. Science, 348(6237):910–914, 2015.

[6] Marco Cuturi. Sinkhorn distances: Lightspeed computation of optimal transport. In Advances in Neural Information Processing Systems, volume 26, 2013.

[7] Pinar Demetci, Rebecca Santorella, Björn Sandstede, William Stafford Noble, and Ritambhara Singh. Gromov-wasserstein optimal transport to align single-cell multi-omics data. bioRxiv, 2020.

[8] Mirjana Efremova and Sarah A. Teichmann. Computational methods for single-cell omics across modalities. Nature Methods, 17(1):14–17, 2020.

[9] Geert-Jan Huizing, Gabriel Peyré, and Laura Cantini. Optimal transport improves cell–cell similarity inference in single-cell omics data. Bioinformatics, 38(8):2169–2177, 2022.

[10] Aditya Kashyap, Maria Anna Rapsomaniki, Vesna Barros, Anna Fomitcheva-Khartchenko, Adriano Luca Martinelli, Antonio Foncubierta Rodriguez, Maria Gabrani, Michal Rosen-Zvi, and Govind Kaigala. Quantification of tumor heterogeneity: from data acquisition to metric generation. Trends in Biotechnology, 40(6):647–676, 2022.

[11] Ilya Korsunsky, Nghia Millard, Jean Fan, Kamil Slowikowski, Fan Zhang, Kevin Wei, Yuriy Baglaenko, Michael Brenner, Po-ru Loh, and Soumya Raychaudhuri. Fast, sensitive and accurate integration of single-cell data with harmony. Nature methods, 16(12):1289–1296, 2019.

[12] Christopher Lance, Malte D. Luecken, Daniel B. Burkhardt, Robrecht Cannoodt, Pia Rautenstrauch, Anna Laddach, Aidyn Ubingazhibov, Zhi-Jie Cao, Kaiwen Deng, Sumeer Khan, Qiao Liu, Nikolay Russkikh, Gleb Ryazantsev, Uwe Ohler, NeurIPS 2021 Multimodal data integration competition participants, Angela Oliveira Pisco, Jonathan Bloom, Smita Krishnaswamy, and Fabian J. Theis. Multimodal single cell data integration challenge: results and lessons learned. Neural Information Processing Systems (NeurIPS) 2021 Competitions and Demonstrations Track, 2022.

[13] Malte Luecken, Daniel Burkhardt, Robrecht Cannoodt, Christopher Lance, Aditi Agrawal, Hananeh Aliee, Ann Chen, Louise Deconinck, Angela Detweiler, Alejandro Granados, Shelly Huynh, Laura Isacco, Yang Kim, Dominik Klein, Bony De Kumar, Sunil Kuppasani, Heiko Lickert, Aaron McGeever, Joaquin Melgarejo, Honey Mekonen, Maurizio Morri, Michaela Müller, Norma Neff, Sheryl Paul, Bastian Rieck, Kaylie Schneider, Scott Steelman, Michael Sterr, Daniel Treacy, Alexander Tong, Alexandra-Chloe Villani, Guilin Wang, Jia Yan, Ce Zhang, Angela Pisco, Smita Krishnaswamy, Fabian Theis, and Jonathan M. Bloom. A sandbox for prediction and integration of DNA, RNA, and proteins in single cells. Proceedings of the Neural Information Processing Systems Track on Datasets and Benchmarks, 1, December 2021.

[14] Noa Moriel, Enes Senel, Nir Friedman, Nikolaus Rajewsky, Nikos Karaiskos, and Mor Nitzan. Novosparc: flexible spatial reconstruction of single-cell gene expression with optimal transport. Nature Protocols, 16(9):4177–4200, 2021.

[15] Aaron van den Oord, Yazhe Li, and Oriol Vinyals. Representation learning with contrastive predictive coding. 1807.03748, 2018.

[16] Adam Paszke, Sam Gross, Francisco Massa, Adam Lerer, James Bradbury, Gregory Chanan, Trevor Killeen, Zeming Lin, Natalia Gimelshein, Luca Antiga, Alban Desmaison, Andreas Köpf, Edward Yang, Zach DeVito, Martin Raison, Alykhan Tejani, Sasank Chilamkurthy, Benoit Steiner, Lu Fang, and Soumith Chintala. Pytorch: An imperative style, high-performance deep learning library. In Neural Information Processing Systems (NeurIPS), volume 32, pages 8024–8035, 2019.

[17] Alec Radford, Jong Wook Kim, Chris Hallacy, Aditya Ramesh, Gabriel Goh, Sandhini Agarwal, Girish Sastry, Amanda Askell, Pamela Mishkin, Jack Clark, Gretchen Krueger, and Ilya Sutskever. Learning transferable visual models from natural language supervision. In International Conference on Machine Learning (ICML), pages 8748–8763, 2021.

[18] Geoffrey Schiebinger, Jian Shu, Marcin Tabaka, Brian Cleary, Vidya Subramanian, Aryeh Solomon, Joshua Gould, Siyan Liu, Stacie Lin, Peter Berube, Lia Lee, Jenny Chen, Justin Brumbaugh, Philippe Rigollet, Konrad Hochedlinger, Rudolf Jaenisch, Aviv Regev, and Eric S. Lander. Optimal-transport analysis of single-cell gene expression identifies developmental trajectories in reprogramming. Cell, 176(4):928–943, 2019.

[19] Tim Stuart and Rahul Satija. Integrative single-cell analysis. Nature Reviews Genetics, 20(5):257–272, 2019.

[20] Alexander Tong, Jessie Huang, Guy Wolf, David Van Dijk, and Smita Krishnaswamy. Trajectorynet: A dynamic optimal transport network for modeling cellular dynamics. In International Conference on Machine Learning (ICML), pages 9526–9536, 2020.

[21] Hongzhi Wen, Jiayuan Ding, Wei Jin, Yiqi Wang, Yuying Xie, and Jiliang Tang. Graph neural networks for multimodal single-cell data integration. In ACM SIGKDD Conference on Knowledge Discovery and Data Mining, pages 4153–4163, 2022.

